# Cell type specific transcriptional reprogramming of maize leaves during *Ustilago maydis* induced tumor formation

**DOI:** 10.1101/552562

**Authors:** Mitzi Villajuana-Bonequi, Alexandra Matei, Corinna Ernst, Asis Hallab, Björn Usadel, Gunther Doehlemann

**Affiliations:** Botanical Institute and Cluster of Excellence on Plant Sciences (CEPLAS), BioCenter, University of Cologne, Zülpicher Str. 47a, Cologne 50674, Germany; Center for Familial Breast and Ovarian Cancer, Medical Faculty, University Hospital Cologne, University of Cologne, Cologne 50931, Germany; BioSC, IBG-2, Institute of Botany, RWTH Aachen, Worringer Weg 3, Aachen 52074, Germany

**Keywords:** *Zea mays*, *Ustilago maydis*, pre-replication complex (pre-RC), SGT1 (suppressor of G2 allele of skp1), endoreduplication, hypertrophy, hyperplasia, tumor

## Abstract

*Ustilago maydis* is a biotrophic pathogen and well-established genetic model to understand the molecular basis of biotrophic interactions. *U. maydis* suppresses plant defense and induces tumors on all aerial parts of its host plant maize. In a previous study we found that *U. maydis* induced leaf tumor formation builds on two major processes: the induction of hypertrophy in the mesophyll and the induction of cell division (hyperplasia) in the bundle sheath. In this study we analyzed the cell-type specific transcriptome of maize leaves 4 days post infection. This analysis allowed identification of key features underlying the hypertrophic and hyperplasic cell identities derived from mesophyll and bundle sheath cells, respectively. We examined the differentially expressed (DE) genes with particular focus on maize cell cycle genes and found that three A-type cyclins, one B-, D- and T-type are upregulated in the hyperplasic tumorous cells, in which the *U. maydis* effector protein See1 promotes cell division. Additionally, most of the proteins involved in the formation of the pre-replication complex (pre-RC, that assure that each daughter cell receives identic DNA copies), the transcription factors E2Fand DPa as well as several D-type cyclins are deregulated in the hypertrophic cells.

## Introduction

*Ustilago maydis* is a biotrophic fungus that triggers tumors in all aerial parts of its host plant maize (*Zea mays*). To attenuate activity of the maize immune system and colonize the different maize organs, *U. maydis* deploys a set of proteins, so called effectors, which manipulate the plant cell metabolism, structure and function for its growth benefit. Such effectors are deployed in a time-, organ- and cell-type-specific manner to reprogram and/or cope with the different maize cell environments^1-11^.

*U. maydis* infection induces characteristic symptoms that include chlorosis, which appears 24 hours post infection (hpi), such lesions are produced in the absence of fungal hyphae suggesting that they result from fungal products such as toxins or effectors^12^. 2 days post infection (dpi) anthocyanin streaking appears and fungal hyphae proliferate and penetrate in between mesophyll cells. At 4 dpi the hyphae have reached the bundle sheath cells and induce tumor formation while at 5 dpi small tumors are visible. 8 dpi maize leaf cells are enlarged and fungal hyphae have undergone branching, a process described as the beginning of teliospore formation^13,14^. Finally, at 12-14 dpi large tumors are formed; inside such tumorous tissue hypha differentiate to give place to the diploid teliospores^15^. Several studies have investigated maize transcriptional reprogramming in response to *U. maydis* infection^10,15–20^. On the cellular level, *U. maydis* induced tumors in maize leaves were found to be constituted of <u>h</u>ypertrophic tumor (HTT) <u>c</u>ells coming from transformed <u>m</u>esophyll cells (M), and <u>h</u>yper<u>p</u>lasic tumor (HPT) cells derived from <u>b</u>undle <u>s</u>heath cells (BS)^4^.

Once induced, maize leaf tumorous cells proliferate even in the absence of the fungus, indicating that *U. maydis* somehow establishes a self-inducing proliferative program in the maize tissues ^21^(Wenzler and Meins, 1986). Remarkably, the cells surrounding the tumors were not able to proliferate, showing that such dedifferentiation and the maintenance of this status is cell-zone specific^21^. Later studies showed that *U. maydis* can extend the undifferentiated state of infected maize tissue^16^. In the leaf this is likely by preventing the establishment of the leaf as a source instead of sink^15,22^. Studies on the maize vascular anatomy and plastid development of intermediate veins show that at the source/sink transition there is minimal development of bundle sheath plastids at the leaf base, as well as in both sections adjoining the source-sink boundary^23^. Therefore successful tumor formation is likely to happen just before the source/sink transition is established suggesting that the “proper” photosynthetic establishment may be crucial to prevent *U. maydis* capacity to induce tumors.

Tumors have been defined as a mass of cells that present abnormal cell divisions and decreased cell differentiation; as a consequence tumors grow in an unorganized way and vary in size and shape ^24^. The cell cycle is tightly regulated and its mechanisms and core machinery are largely conserved among eukaryotes^25-27^. Two key regulatory molecules determine cell cycle progression; cyclins and cyclin-dependent kinases (CDKs)^26^. CDKs are known as master cell cycle regulators and must associate with their regulatory cyclin partner to be active^26^. Besides, CDK activity is regulated in other ways including changes in the phosphorylated status, interaction with inhibitory proteins or non-catalytic CDK-specific inhibitors (CKIs), and proteolysis by the 26S proteosome^28,29^. Two major classes of CDKs can be distinguished, CDKA and CDKB^26^. CDKA regulate the G1-to-S and G2-to-M-transitions while CDKB control the G2-to-M transition^26^. Plants encode for cyclins grouped as A-, B-, and D-types^26^. A-type cyclins control mainly S-phase and the G2/M transitions; B-type cyclins control G2/M transition, while D-type cyclins are involved in G1/S transition^28,30^. Two major multimeric E3 ubiquitin ligases target cell cycle regulators to the proteasome to promote cell cycle progression: the anaphase promoting complex/cyclosome (APC/C) and Skp1/Cullin/F-box complex^29,31^. APC/C is multiprotein complex and controls the exit from mitosis by targeting important mitotic promoting proteins like cyclin B for degradation via the 26S proteasome^29^. SCF regulates mainly the G1-to-S transition by degrading CDK inhibitors (CKIs) like ICK/KRP proteins^31,32^. The cell cycle is relatively well functionally characterized in the plant model *Arabidopsis thaliana*; in contrast, less is known about the roles of key cell cycle-controlling genes in maize^33,34^.

This study is combining the high-resolution technique Laser Capture Microdissection with high transcriptome profiling RNAseq to characterize maize tumorous mesophyll and bundle sheath cells induced by *U. maydis* infection. In a previous article we have described the *U. maydis* transcriptome showing the specificity of effector deployment in a cell type-specific manner^4^. We now describe the maize-specific transcriptome response of micro-dissected mesophyll and bundle sheath tumorous cells. Moreover, we take the information of an *U. maydis* effector deletion mutant, SG200Δsee1^8^, which induces hypertrophic but not hyperplasic tumors in maize leaves after infection^4^ to pinpoint possible cell-cycle related genes and/or the mechanism that could explain the observed phenotype. Since tumors are a product of cell cycle alterations, we analyze this cellular process in a deeper detail.

## Materials and Methods

Plant growth conditions, fungal infections, tissue embedding, sectioning, single-cell LCM and RNA sequencing details are fully described in Matei et al., 2018.

### RNAseq analysis

The quant utility of kallisto v0.43.1^35^ was used for alignment-free estimation of RNAseq read abundancies in a merged reference genome consisting of Zea *mays* B73 RefGen_v3^36^ and Ustilago *maydis* 521 v2.0 available at NCBI Genomes Server (ftp://ftp.ncbi.nih.gov/genomes/), using the supplied annotations. Resulting estimated counts served as input for differential expression tests with sleuth v0.29.0^37^ following the protocol described at https://pachterlab.github.io/sleuth_walkthroughs/trapnell/analysis.html. Differential expressed (DE) genes were selected due to criterion p-value <0.05 after correction of p-values for multiplicity using the Benjamini-Hochberg approach with FDR set to 0.05^38^.

Expression changes were assessed as Log2Fold-change calculated as cell-type specific infected (treated) divided by the cell-type specific uninfected (untreated) values. The experimental design allowed six comparisons: mock bundle sheath cells against mock mesophyll cells (MBS.vs.MMS), mock bundle sheath cells against SG200 infected bundle sheath cells (MBS.vs.HPT), mock mesophyll cells against SG200 infected mesophyll cells (MMS.vs.HTT), SG200Δsee1 infected mesophyll cells against mock mesophyll cells (MMS.vs.seeTC), SG200Δsee1 infected mesophyll cells against SG200 infected mesophyll cells (SeeTC.vs.HTT) and SG200 infected bundle sheath cells against SG200 infected mesophyll cells (HPT.vs.HTT).

### Core Cell Cycle Gene List

This table was generated based on the MapMan Bin annotations using Mercator4 V1 and annotating data reported in the literature. A full table including all cell cycle related processes like: organelle machinery for DNA replication and organelle fission, cytokinesis, chromatin condensation, sister chromatid separation, chromosome segregation and DNA damage response is provided in the supplementary material (Supplementary Table 3).

### Core DNA Replication Machinery Gene List

This list is based on the published table by Shultz et al., 2007. We look for the reported orthologues and functional annotations provided at Joint Genome Institute (JGI), Zmays, 5b+, annotation, file: Zmays: Zmays_284_5b+.annotation_info.txt. Based on that annotation we seek for Arabidopsis homologues and introduced the reported maize gene. To keep the coherence with the table reported by Shultz et al., 2007, Rice locus homologues were searched for syntenic orthologues provided at Freeling lab (ftp.maizegdb.org/FTP/bulk/grass_syntenic_orthologs.csv), which are mapped to RefGen_v2. In general, by this method we detect different maize putative homologues per Arabidopsis gene. To facilitate table read the reported orthologues are bold letters prioritizing the orthologues between maize and rice (Supplementary Table 4).

### Skp1/Cullin1/F-box complex (SCF) Genes Table

This table was generated based on the MapMan Bin annotations using Mercator4 V1. In addition LRR-maize genes were annotated based on the data provided by Song et al., 2015.

### Small ubiquitin-like modifier (SUMO) and the SUMOylation Machinery Genes Table

This table was generated based on the information provided by Augustine et al., 2016, in this paper a full analysis and description of the SUMO system in maize has being thoroughly performed by these topic experts. Lectors interested on the topic please refer to that publication (Supplementary Table 5).

### Gene Ontology (GO) and metabolic pathway analyses

Differentially expressed (DE) genes were analyzed for gene ontology (GO) enrichment with the web-based agriGO software^39^. To identify possible connections among DE genes list we applied the Batch SEA function SEACOMPARE, which allows comparisons among the significant GO terms from the results of selected datasets to effectively identify GO terms.

## RESULTS

### Analysis of the maize transcriptional response during tumor formation in hyperplasic and hypertrophic cells

In a previous work we showed that *U. maydis*-induced maize tumors are constituted of hypertrophic mesophyll tumor (HTT) cells coming from the mesophyll cells and small hyperplasic tumor (HPT) cells coming from the bundle sheath cells^4^. In contrast to the solopathogenic strain SG200^3^, the *U. maydis* effector deletion mutant SG200Δsee1 fails to induce DNA synthesis and cell division in bundle sheath cells^4^. Consequently, tumors caused by SG200Δsee1 are mainly built of HTT; while the bundle-sheath derived small tumor cells are missing^4^. To determine genes differentially expressed (DE) in each particular tumorous cell type we performed six pairwise comparisons of cell-type specific control mock groups against SG200 or SG200Δsee1 infected cells (Table I; Supplementary Dataset 1). The highest number of DE genes was observed in the SG200 infected cells (Table I), either HTT or HPT when compared to uninfected/mock cells (Mock Bundle Sheath, MBS; Mock Mesophyll cells, MMS), indicating that *U. maydis* infection induces a strong transcriptional maize cell reprogramming, which is in agreement with previous reports^15,20^. SG200Δsee1 infection induces a milder effect in the mesophyll cells; we found less DE genes when we compared SG200Δsee1 to the MMS than to the HTT (Table I). This is in agreement with the mild SG200Δsee1 tumor phenotype observed, where the bundle sheath structure is largely preserved and the hypertrophic cells are mostly absent^4^. Few DE genes were detected when we compared HTT against HPT, suggesting that many of the DE genes are shared between these two datasets and their expression behaviors are likely similar (Table I).

More genes are up-regulated than down-regulated in response to SG200 infection in both cell types (Figure 1A). The largest difference is observed in the HTT dataset where 6,852 are upregulated in contrast to 1,504 downregulated genes (Figure 1A). To determine a significant change in gene expression we applied an arbitrary absolute log2-Fold Change (log2FC) threshold of 1.5. This cutoff drastically reduced the number of DE genes; however it kept the observed tendency of more genes being upregulated than downregulated (Figure 1A).

**Figure 1.**
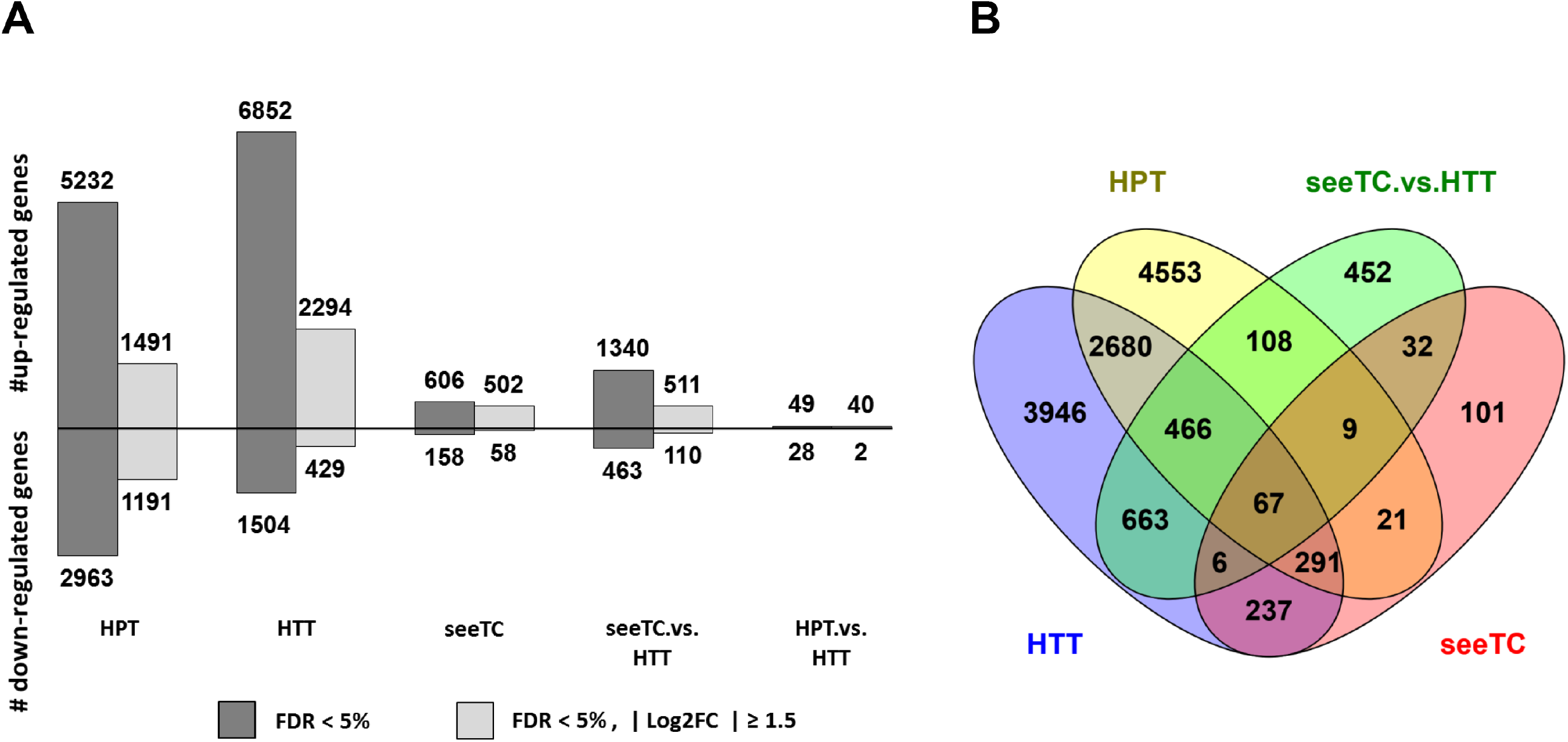
RNA sequencing profiling identifies DE genes for each cell-type specific tumor. A) Number of differentially expressed genes per dataset. Bars represent up and down-regulated genes in the five pairwise comparisons of cell-type specific mocked groups against infected. For the last two datasets, seeTC.vs.HTT and HPT.vs.HTT, HTT was defined as the control group. B) Venn diagram showing the number of shared and unique DE genes revealed by pairwise comparison^103^.

A small number of genes is DE in all considered datasets (67 genes); in contrast, many genes are shared between HTT and HPT datasets (2680 genes, Figure 1B). HPT contains the highest number of uniquely expressed genes (4553), followed by HTT (3946), and seeTC (101, Figure 1B).

In summary, the results demonstrate that SG200 infection has a strong effect on gene expression in both mesophyll and bundle sheath cells. This is in line with the observation that tumor formation correlates with a strong cell reprogramming^15^ and this may involve the gene expression of otherwise silenced genes. Moreover, the See1 effector seems to have a key role in such response since the number of DE genes is drastically reduced in SG200Δsee1 infected mesophyll cells (seeTC) in comparison to SG200 infected cells (HTT, Figure 1B). This gene expression profile reflects the phenotype, as SG200Δsee1 infections induce small tumors^4,8^.

### Functional categorization of DE genes in the hyperplasic and hypertrophic cells: Gene Ontology enrichment (GO) analysis

To explore the nature of the data we analyzed all DE genes for Gene Ontology enrichment (GO) with the web-based agriGO software^39^. The Singular Enrichment Analysis (SEA) revealed a strong and shared enrichment for several GO-terms between HPT and HTT datasets (Supplementary Dataset 2 and Supplementary Table 1). A total of 171 different GO terms were assigned to all five datasets. 101 terms are common between HPT and HTT datasets. In contrast, only 23 GO terms are common between HTT and seeTC datasets, which is interesting as the only difference is the deletion of the See1 effector supporting the strong effect of this protein. We further analyzed the data with Parametric Analysis of Gene set Enrichment (PAGE), which takes the expression levels into account. This analysis showed 163, 77 and 15 GO terms for HPT, HTT and seeTC respectively (Supplementary Dataset 2). The majority of the GO terms found in HTT and seeTC datasets are shared with HPT with exception of 13 unique to HTT and 4 unique to seeTC datasets. These include very diverse functions in HTT and kinase and transferase activities for seeTC (Supplementary Dataset 2 and Supplementary Table 2).

**Table I.** Differentially expressed (DE) genes in the six datasets

| Comparison - Dataset | DE genes full<sup>a</sup> | DE genes unique<sup>b</sup> |
| --- | --- | --- |
| MBS.vs.HPT | 8 195 | 4 553 |
| MMS.vs.HTT | 8 356 | 3 946 |
| MMS.vs.seeTC | 764 | 101 |
| seeTC.vs.HTT | 1 803 | 452 |
| HPT.vs.HTT | 77 | 6 |

<sup>a</sup> DE genes full contain all genes after applying a threshold of $|\log_2\text{FC}| \geq 1.5$ .
<sup>b</sup> DE genes unique indicates the number of DE genes ( $|\log_2\text{FC}| \geq 1.5$ ) that are exclusive for the corresponding dataset.

**Table II.**
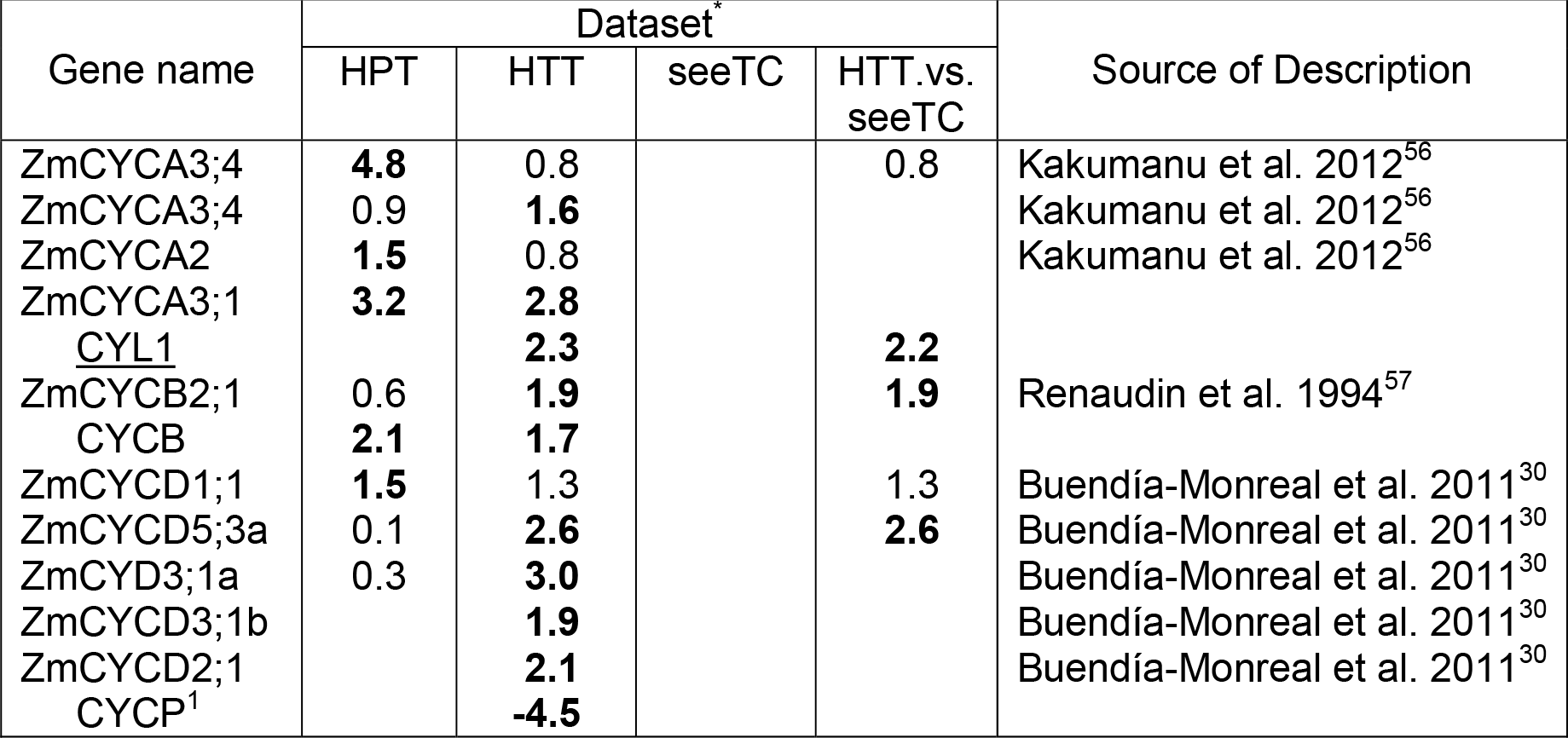
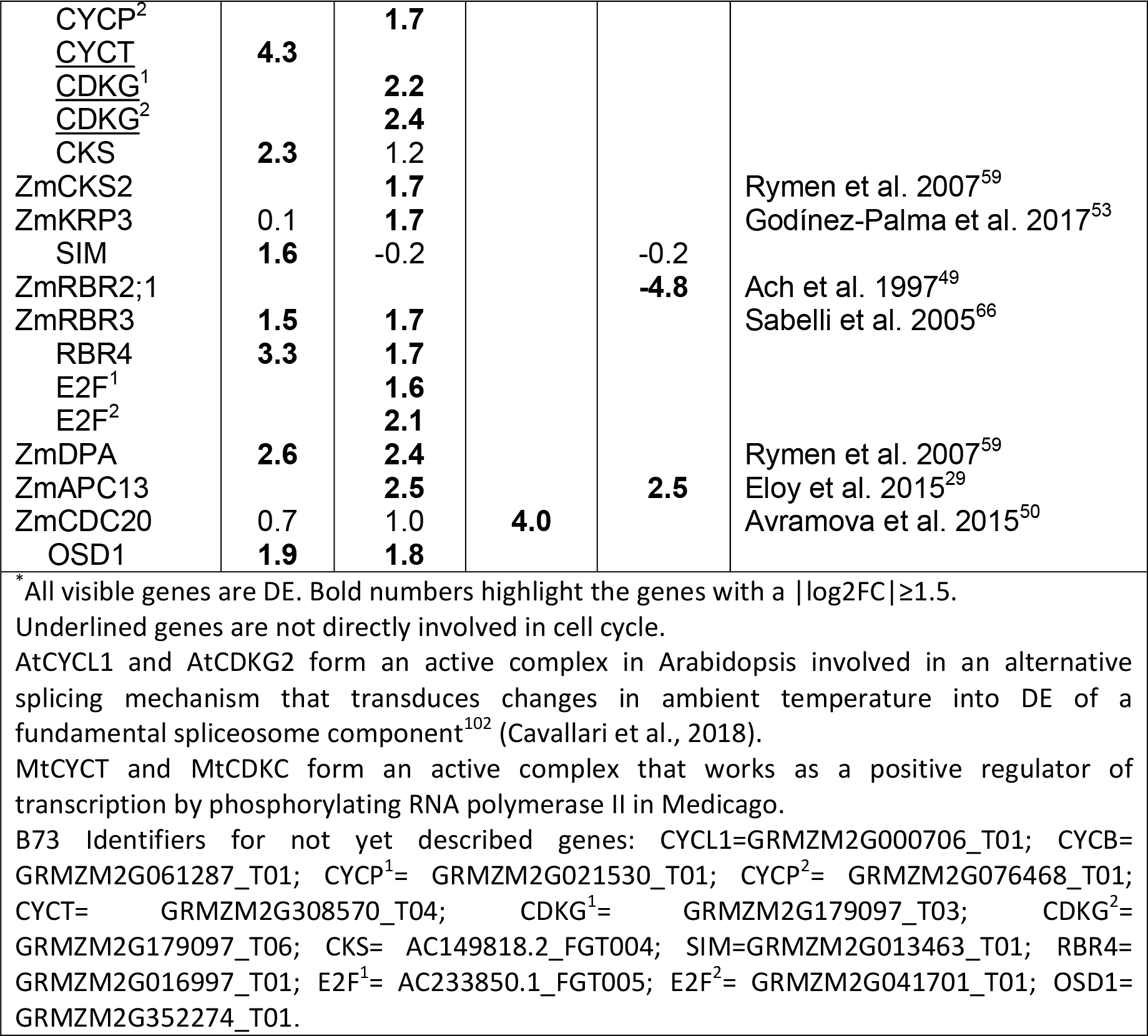
DE basic cell cycle machinery genes in the different datasets

| Gene name | Dataset |  |  |  | Source of Description |
| --- | --- | --- | --- | --- | --- |
|  | HPT | HTT | seeTC | HTT.vs.seeTC |  |
| ZmCYCA3;4 | **4.8** | 0.8 |  | 0.8 | Kakumanu et al. 2012<sup>56</sup> |
| ZmCYCA3;4 | 0.9 | **1.6** |  |  | Kakumanu et al. 2012<sup>56</sup> |
| ZmCYCA2 | **1.5** | 0.8 |  |  | Kakumanu et al. 2012<sup>56</sup> |
| ZmCYCA3;1 | **3.2** | **2.8** |  |  |  |
| CYL1 |  | **2.3** |  | **2.2** |  |
| ZmCYCB2;1 | 0.6 | **1.9** |  | **1.9** | Renaudin et al. 1994<sup>57</sup> |
| CYCB | **2.1** | **1.7** |  |  |  |
| ZmCYCD1;1 | **1.5** | 1.3 |  | 1.3 | Buendía-Monreal et al. 2011<sup>30</sup> |
| ZmCYCD5;3a | 0.1 | **2.6** |  | **2.6** | Buendía-Monreal et al. 2011<sup>30</sup> |
| ZmCYD3;1a | 0.3 | **3.0** |  |  | Buendía-Monreal et al. 2011<sup>30</sup> |
| ZmCYCD3;1b |  | **1.9** |  |  | Buendía-Monreal et al. 2011<sup>30</sup> |
| ZmCYCD2;1 |  | **2.1** |  |  | Buendía-Monreal et al. 2011<sup>30</sup> |
| CYCP<sup>1</sup> |  | **-4.5** |  |  |  |
| CYCP<sup>2</sup> |  | **1.7** |  |  |  |
| <u>CYCT</u> | **4.3** |  |  |  |  |
| <u>CDKG<sup>1</sup></u> |  | **2.2** |  |  |  |
| <u>CDKG<sup>2</sup></u> |  | **2.4** |  |  |  |
| CKS | **2.3** | 1.2 |  |  |  |
| ZmCKS2 |  | **1.7** |  |  | Rymen et al. 2007<sup>59</sup> |
| ZmKRP3 | 0.1 | **1.7** |  |  | Godínez-Palma et al. 2017<sup>53</sup> |
| SIM | **1.6** | -0.2 |  | -0.2 |  |
| ZmRBR2;1 |  |  |  | **-4.8** | Ach et al. 1997<sup>49</sup> |
| ZmRBR3 | **1.5** | **1.7** |  |  | Sabelli et al. 2005<sup>66</sup> |
| RBR4 | **3.3** | **1.7** |  |  |  |
| E2F<sup>1</sup> |  | **1.6** |  |  |  |
| E2F<sup>2</sup> |  | **2.1** |  |  |  |
| ZmDPA | **2.6** | **2.4** |  |  | Rymen et al. 2007<sup>59</sup> |
| ZmAPC13 |  | **2.5** |  | **2.5** | Eloy et al. 2015<sup>29</sup> |
| ZmCDC20 | 0.7 | 1.0 | **4.0** |  | Avramova et al. 2015<sup>50</sup> |
| OSD1 | **1.9** | **1.8** |  |  |  |

\* All visible genes are DE. Bold numbers highlight the genes with a $|\log_2FC| \geq 1.5$ . Underlined genes are not directly involved in cell cycle. AtCYCL1 and AtCDKG2 form an active complex in Arabidopsis involved in an alternative splicing mechanism that transduces changes in ambient temperature into DE of a fundamental spliceosome component<sup>102</sup> (Cavallari et al., 2018). MtCYCT and MtCDKC form an active complex that works as a positive regulator of transcription by phosphorylating RNA polymerase II in Medicago. B73 Identifiers for not yet described genes: CYCL1=GRMZM2G000706\_T01; CYCB=GRMZM2G061287\_T01; CYCP<sup>1</sup>= GRMZM2G021530\_T01; CYCP<sup>2</sup>= GRMZM2G076468\_T01; CYCT= GRMZM2G308570\_T04; CDKG<sup>1</sup>= GRMZM2G179097\_T03; CDKG<sup>2</sup>= GRMZM2G179097\_T06; CKS= AC149818.2\_FGT004; SIM=GRMZM2G013463\_T01; RBR4= GRMZM2G016997\_T01; E2F<sup>1</sup>= AC233850.1\_FGT005; E2F<sup>2</sup>= GRMZM2G041701\_T01; OSD1= GRMZM2G352274\_T01.

**Table III.**
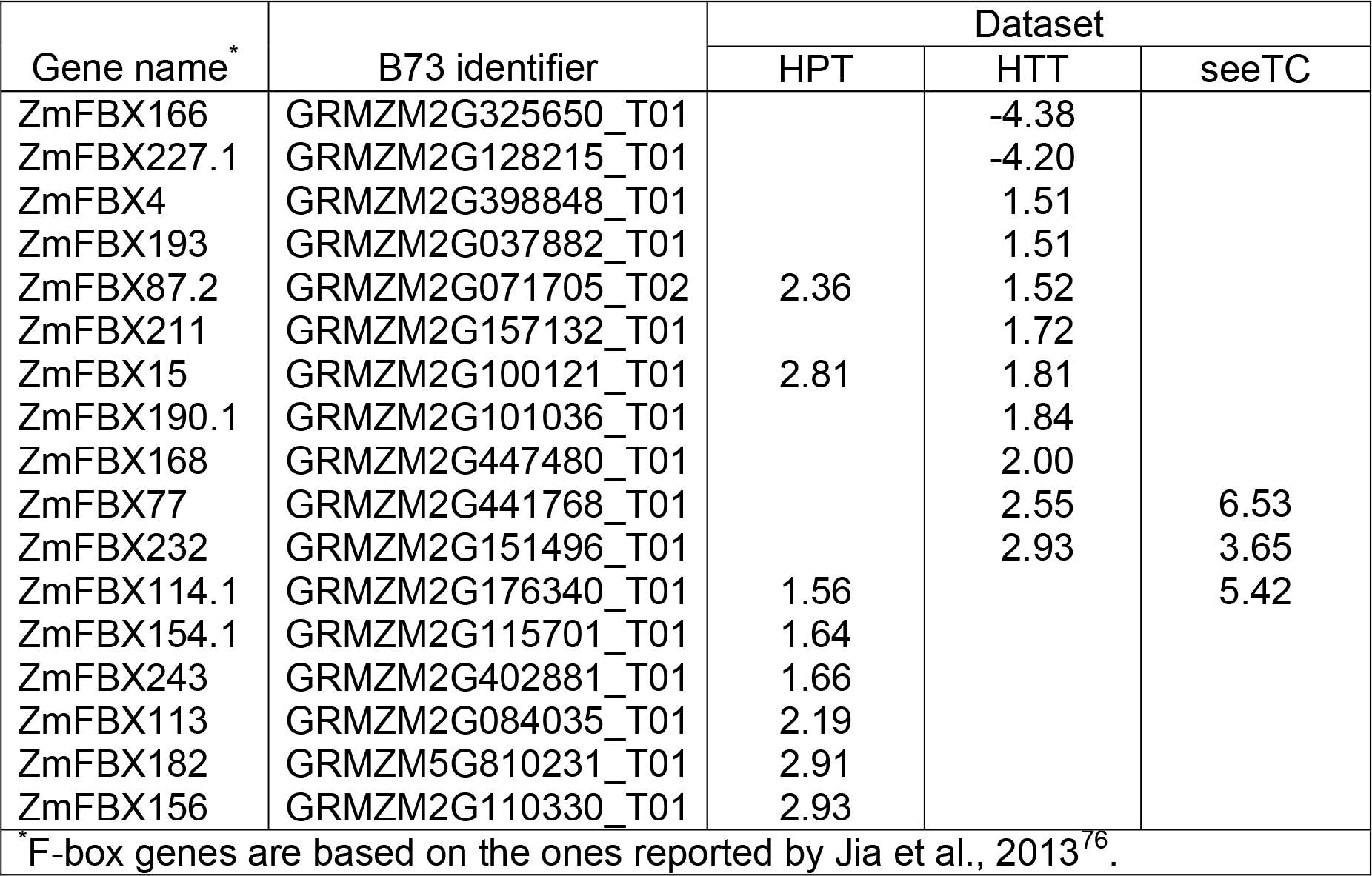
DE F-box genes in the different datasets

Since a considerable number of genes were DE genes and unique in each dataset (Table I and Figure 1B), we decided to explore if such gene subsets were also enriched for specific GO terms. After SEA analysis we found 55 HPT and 44 HTT GO enriched terms, out of which 31 are shared (Supplementary Dataset 3). This suggests that similar functions are performed by different genes that are expressed specifically in each cell-type. Also interesting is that fewer GO terms are lost in the HTT dataset, when comparing the terms assigned to the full list of DE genes against unique DE genes (77 full vs. 44 unique), than in the HPT dataset (163 full vs. 55 unique), suggesting that a lot of the functional diversity for HTT is contain/shared within the unique DE genes. Further analysis with PAGE showed 40 GO terms enriched for HPT and only 5 for HTT (Supplementary Dataset 3). These last five terms are shared with HPT dataset and include: GO:0010467-gene expression; GO:0034645-cellular macromolecule biosynthetic process; GO:0009059-macromolecule biosynthetic process; GO:0043229-intracellular organelle and GO:0043226–organelle. The remaining datasets showed no enrichment (Table I).

### Dissecting of differentially regulated biological processes: MapMan-Bin enrichment analysis

For a more detailed and less redundant functional classification of DE genes a MapMan-Bin enrichment analysis was performed. MapMan is a software tool composed of different modules including a set of Scavangers which assign non-redundant functional categories to a set of given genes, proteins or metabolites and an Image Annotator module which allows the visualization of data on diagrams of biological processes or pathways relying on mapping files created by the scavangers^40,41^. The plant gene function ontology MapMan consist of 34 major bins and is organized as a tree, thus enabling the categorization of gene functions at different levels of generality^41^. Here, we used direct (level one) children of the root node to generate the profiles, counting all annotations by their respective level one. Afterwards we tested for overrepresented terms in the intersection (both terms), difference (one but not the other) and union (either) in mesophyll and bundle sheath datasets infected with SG200 using exact Fischer tests^42^. This analysis showed an overrepresentation of five MapMan-Bins (Figure 2A). Terms include chromatin assembly and remodeling (histones, H4-type histone - 12.1.5), cell-cycle (regulation cyclins, CYCA-type cyclin – 13.1.1.1), protein biosynthesis (translation and elongation, eEF1A aminoacyl-tRNA binding factor – 17.4.1), cytoskeleton (microtubular network, kinesin microtubule-based motor protein activities, kinesin-5 motor protein – 20.1.3.4) and protein modification (phosphorylation, TKL kinase superfamily - 18.9.1). Interestingly, when looking for the expression status of genes annotated with any of these five MapMan-Bins we find genes that are both DE and tissue specific (Figure 2B). This indicates that while the respective gene functions (MapMan-Bins) are shared and characteristic for both tumor tissues, there are tissue specific DE genes implementing these functions. Based on these results, and the clear implication of the deregulation of cell cycle regulating genes in tumor formation, we further examined the DE genes with particular focus on maize cell cycle genes.

**Figure 2.**
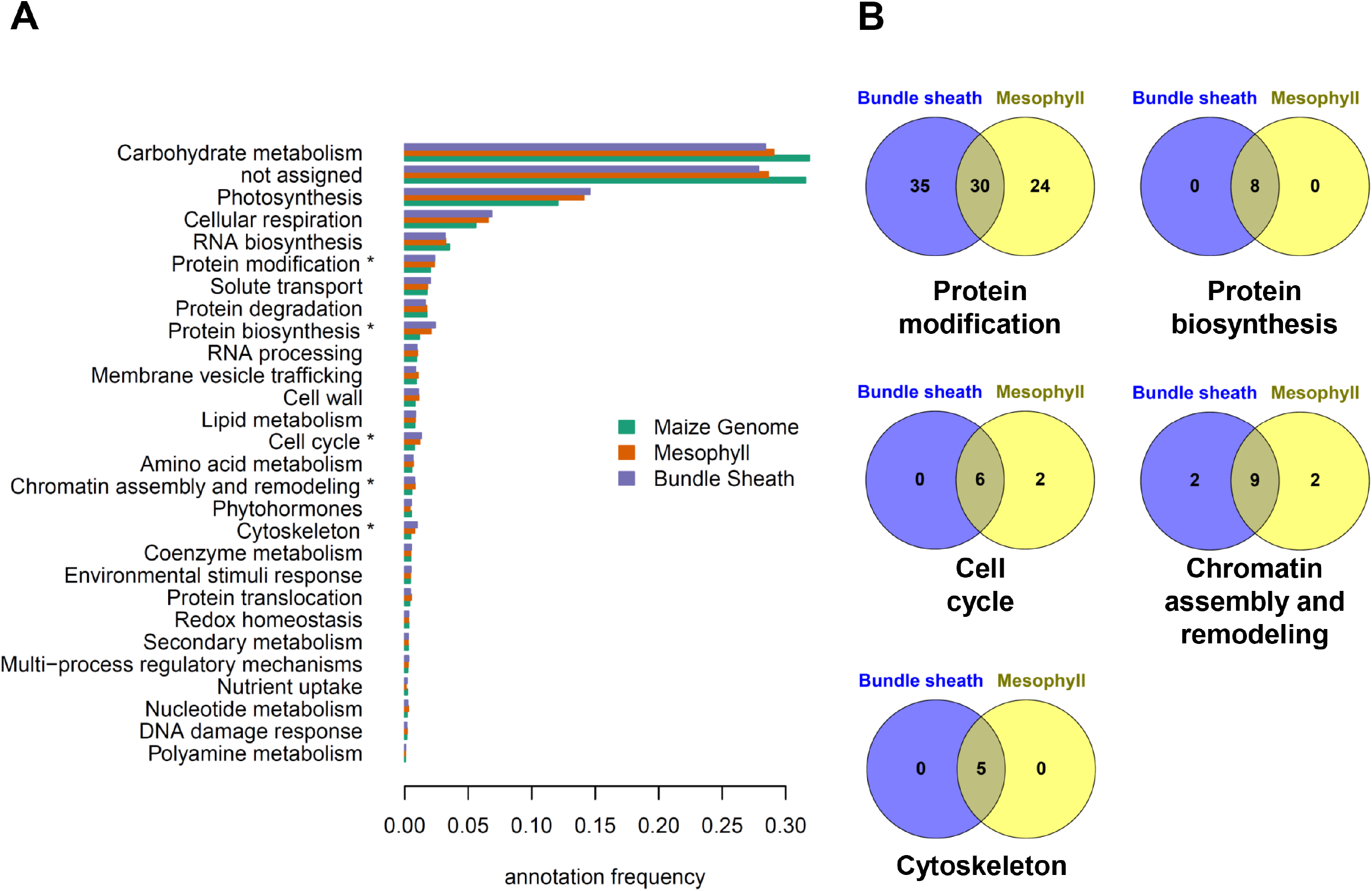
Overrepresentation analyses of DE genes within functional MapMan-Bins categories. A) Gene function profile of the maize genome (green) compared to the two *U. maydis* infected maize tissues: mesophyll (orange) and bundle sheath (violet). Profiles were generated using the 35 major MapMan-bin annotations, from which 28 are shown. P-Values resulting from exact Fischer tests were corrected for multiple hypothesis testing using false discovery rate estimates. Asterisk denotes the enriched bin categories. B) Venn diagrams for the five significant MapMan-bins identified. DE genes are indicated with numbers and percentage in brackets.

For a general overview of the effect of *U. maydis* infection in maize mesophyll and bundle sheath cells we generate a metabolic overview map with MapMan^40,41^. The strongest effect is observed in the photosynthetic light reactions section for the HPT dataset, where many genes are downregulated (Figure 3 and Supplemental Figure 1). Comparably, genes involved in starch formation were downregulated in the HPT dataset (Figure 3 and Supplemental Figure 2). This is in agreement with our previous finding that these cells are depleted from chloroplasts^4^. In contrast, starch formation and degradation related genes were slightly but mostly upregulated in the HTT dataset (Figure 3 and Supplemental Figure 2). This provides a picture of the maize leaf response towards *U. maydis* infection and supports the hypothesis of HPT working as a strong active sink tissue that stimulates the attraction of nutrient flow from source tissues, which in this case might be partially enabled by HTT^4,43^. For the seeTC dataset we observe mostly strong upregulation in very punctual but overall distributed processes (Figure 3).

**Figure 3.**
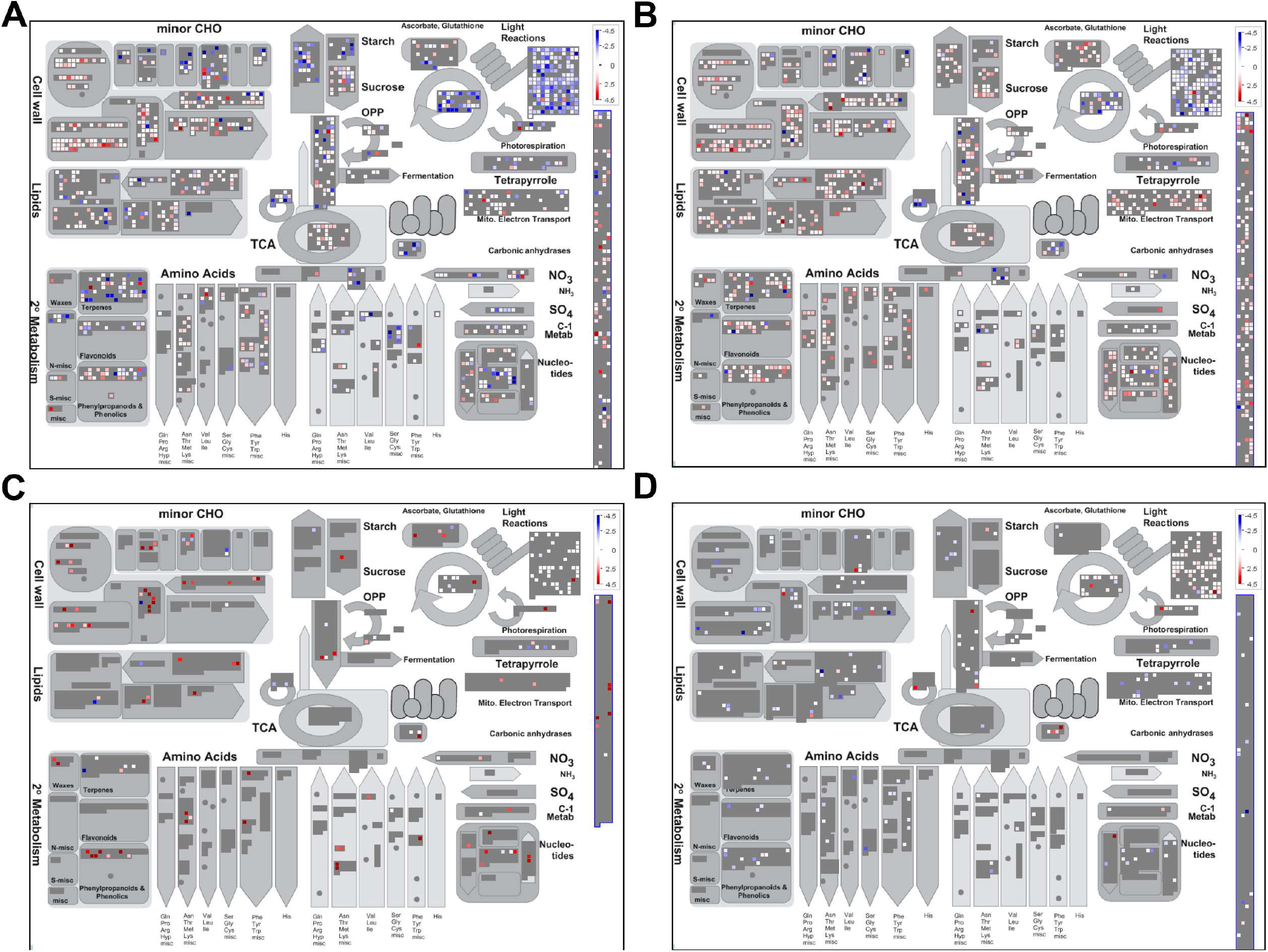
Overview of maize metabolic responses to *Ustilago maydis* infection in specific cell-types. Genes differentially expressed (FDR ≤5%) are shown **A**, HPT: Mapped 23299 out of 28034 (some of the data points may be mapped multiple times to different bins). Visible in this pathway: 2818. **B** HTT: Mapped 22714 out of 27240 (some of the data points may be mapped multiple times to different bins). Visible in this pathway: 2835. **C** seeTC: Mapped 14429 out of 17311 (some of the data points may be mapped multiple times to different bins). Visible in this pathway: 1901. **D** seeTC.vs.HTT: Mapped 24454 out of 29330 (some of the data points may be mapped multiple times to different bins). Visible in this pathway: 3040. Upregulated transcripts are shown in red and downregulated transcripts are colored blue.

In maize, cellulose microfibrils are mainly crosslinked with glucuronoarabinoxylans (GAXs)^44^. During cell elongation in growing tissues the mixed-linkage (1→3), (1→4)- *β*-d-glucan appears transiently as the major cross-linking glycan^45^. Analysis of gene expression of cell wall precursors in HPT and HTT datasets show an upregulation for genes involved in the transformation from UDP-D-glucose to: sucrose, UDP-L-rhamnose, UDP-D-galacturonic acid and UDP-D-xylose (Supplemental Figure 3). Interestingly, the conversion of UDP-D-xylose to UDP-L-arabinose is upregulated in the HPT dataset (Supplemental Figure 3). This is in agreement with our data which indicate that *U. maydis* infection change the ratio contents of monosaccharides, increasing arabinose content and reducing xylose ^4^.

We have previously shown that tumors develop and expand in between two primary leaf veins where lignin deposition increases defining the tumor borders^4^. Lignification is commonly associated with plant defense response. The HTT dataset shows an upregulation of genes involved in the formation of three important lignin precursors, namely p-coumaryl alcohol, coniferyl alcohol and sinapyl alcohol^46,46^ (Supplemental Figure 4).

### Tissue-specific regulation of cell-cycle associated genes by *U. maydis*

The See1 effector is required for the activation of maize cell mitotic division in bundle sheath cells^4,4^. Therefore, we asked if the unique subset of maize cell cycle genes DE in the bundle sheath SG200 infected cells could reflect the processes that are likely See1-driven.

Cell cycle comprises a sequence of events including DNA replication, cell division and growth all of them requiring the precise coordination of several protein complexes^48^. A candidate list of core-cell-cycle genes was generated using the MapMan Bin annotations using Mercator4 V1 and edited based on literature search to include/annotate described maize cell-cycle machinery and core regulator genes^25,29,30,49–61^ (Supplementary Table 3). To facilitate the analysis the core DNA replication machinery (pre-replication complex and genes involved in s-phase) is analyzed in the next chapter. In general, genes that constitute the basic cell cycle machinery appear DE in four out of five datasets after setting a threshold of |log2FC| ≥1.5 (Table II). The HTT dataset present the highest number of DE genes (22), followed by HPT (12), HTT.vs.seeTC (5) and seeTC (1; Table II). The maize genome encodes over 50 cyclins, the majority of which remain uncharacterized^62,63^. Three cell cycle related cyclins, namely A-, B- and D-types are DE in the HPT and HTT datasets. A-type cyclins, which normally are involved in S-phase progression, are upregulated in both HPT and HTT datasets. *GRMZM2G017081*, which encodes for an A2-type cyclin, is upregulated in the HPT dataset, and was found as part of a subnetwork that positively correlates with leaf size and timing traits in maize^64^. Two uncharacterized B-type cyclins appear upregulated in the HPT and HTT datasets. B-type cyclins are key actors in the G2/M transition and expressed in a narrow time window from late to mid M phase^62^. Finally, several D-type cyclins are upregulated in the HTT dataset. D-type cyclins are regarded as G1-specific and proposed to be sensors of growth conditions by integrating internal and environmental cues^48,65^. Particularly, ZmCYCD2;1 has a positive role in the endoreduplication cycle in endosperm^33^.

Maize contains at least four Retinoblastoma-related (*RBR)* genes that can be functionally grouped as repressors *RBR1/2* and promoters *RBR3/4* of the E2F-DP factors, which promote the transcription of genes required for cell cycle progression^34,66–68^. We observe an up-regulation of *RBR3/4* genes in the HPT and HTT cells. Additionally *E2F/DP* coding genes are upregulated in HTT cells (Table II).

APC13 is upregulated in the HTT dataset. In humans and yeast APC13 is required for efficient cyclin degradation by promoting the association of the APC3 and APC6 subunits, until now APC13 has not been characterized in plants^29^. CDC20 is strongly upregulated in seeTC dataset. CDC20 is a crucial co-activator of APC/C to degrade Securin and CYCB, promoting in this way the onset of anaphase and mitotic exit^29^.

*OMISSION OF SECON DIVISION 1 (OSD1)/ GIGAS CELL 1(GIG1)* expression levels peak at the G2/M transition. Arabidopsis plants overexpressing *OSD1/GIG1* accumulate CYCB1;2^29^. We detect an upregulation of two B-type cyclins in the HPT and HTT cells suggesting a similar effect (Figure 4).

**Figure 4.**
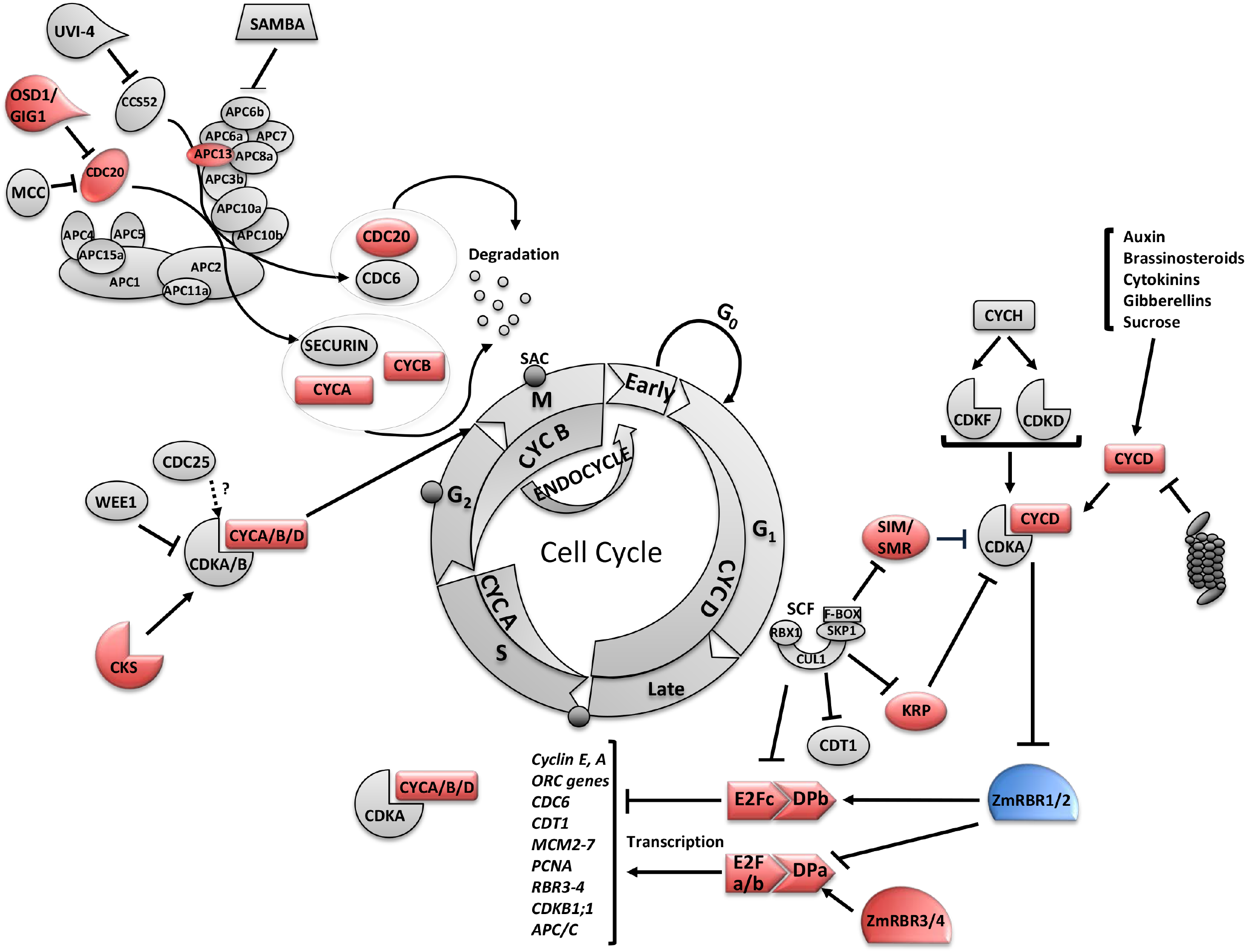
General cell-cycle model highlighting the genes that respond to *Ustilago maydis* infection. Cell cycle consist of four phases: G1, S, G2 and M. S-phase (Synthesis) is where DNA replication takes place, and M-phase (Mitosis), where nuclear division occurs. G1 and G2 are gap phases where some checkpoints to control cell cycle progression take place. A more detailed explanation of the cycle is given in the text. Genes differentially regulated (|log2FC|≥1.5) are shown. Circles at the beginning of S-phase and middle of G2 and mitosis represent cell-cycle progression checkpoints. Colors represent gene expression profile: red = upregulated and blue = downregulated.

Two CDK subunit (CKS) proteins are upregulated in the HPT and HTT datasets. CKS work as scaffold proteins that serve as adaptors for targeting CDKS to mitotic substrates but in contrast to cyclins, are not required for proper phosphorylation activity^69-71^.

Two CDK Inhibitors (CKIs), belonging to different groups are DE in the HPT and HTT datasets. GRMZMM2G013463, which encodes for an uncharacterized SIAMESE gene (SIM) is upregulated in HPT, and a Kip-Related protein (KRP) ZmKRP3, which is upregulated in HTT dataset. ZmKRP3 belongs to a class of KRPs exclusively present in monocotyledonous plants and presents motifs required for the interaction with CDKs and D-type cyclins, shows a PEST sequence required for targeted degradation and does not present a nuclear localization signal^53^.

In Arabidopsis, there is a concentration-dependent role of ICK/KRPs in blocking both the G1/S cell cycle and entry into mitosis but allowing S-phase progression promoting a switch to endoreduplication^72^. Several of the best-characterized SIAMESE (SIM) and SIAMESE RELATED (SMR) proteins are also involved in the regulation of the transition from the mitotic cell cycle to endoreplication^72,73^. This poses the question if the two distinct CKI upregulated in the different tumorous cell types are inducing different outcomes to give place to hyperplasic or hypertrophic phenotypes. At least our data clearly indicate that nuclear size, which can be proportionally related to endoreduplication, of mesophyll cells infected with SG200 or SG200Δsee1 is increased while bundle sheath nuclear sizes remain unchanged (Figure 5). This supports the concept of hypertrophy in mesophylls cells being linked to endoreduplication.

**Figure 5:**
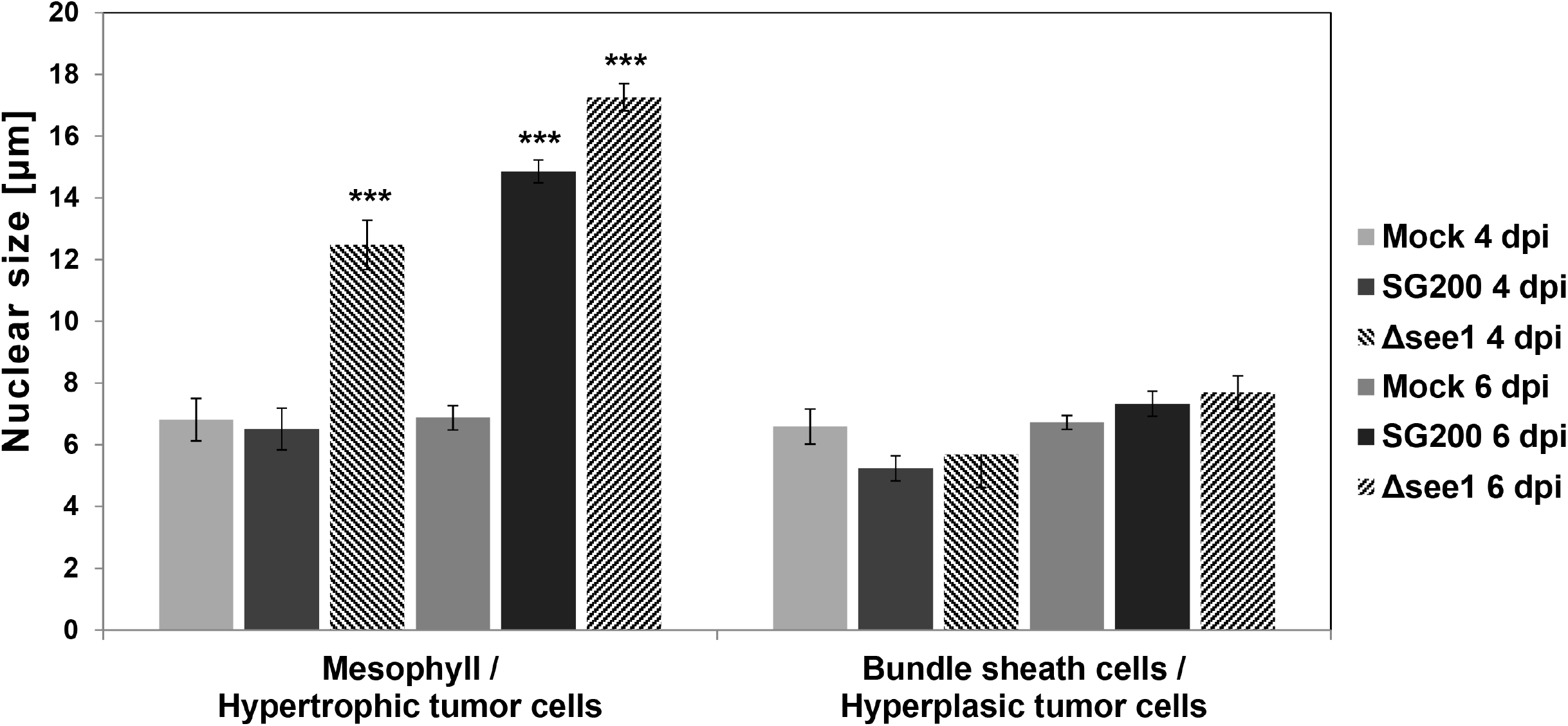
Cell type-specific nuclear size measurements in leaf tissue sections stained with propidium iodide (PI) at major tumor development stages. Data shows nuclear size measurements 4 dpi and 6 dpi in mock treated, SG200 infected and SG200Δsee1 infected PI stained leaf tissue sections. Analyzed cell types included Mesophyll and resulting hypertrophic tumor cells, as well as bundle sheath cells and resulting hyperplasic tumor cells. A minimum of 70 nuclei was measured per tissue type. Results represent the mean ± SD from three independent leaf sections per biological replicate. Two independent biological experiments were performed. Asterisks indicate statistical significance of nuclear size compared to mock treated tissue of the same age. P-values were calculated using the unpaired student’s t-test; ***: p≤ 0.001.

### The Pre-Replication complex (pre-RC, before S-phase)

The pre-RC is a very important part of the cell cycle as it defines the origins to initiate DNA replication, regulates DNA replication and assures that each daughter cell receives identic DNA copies^51^. Pre-RC members are conserved in all eukaryotes and previous studies have shown that plants core DNA replication machinery is more similar to vertebrates than single celled yeasts^25-27^. The pre-RC consist of an initiator to establish the site of replication initiation (ORC), a helicase to unwind DNA (MCM complex), and CDC6 and CDT1, which act synergistically to load the MCM complex^25,51^. The formation of a pre-replication complex (pre-RC) is a key control mechanism occurring before cells enter S-phase^51^.

Our analysis shows that DE genes from the core DNA replication genes are found in three of the five datasets (Figures 6 and 7 and Supplementary Table 4), the HPT dataset (11 genes), HTT (22 genes), and SeeTC.vs.HTT (3 genes). In the HPT dataset we found exclusively upregulated ORC5, ORC6, CDC6, CDT1 (b), SLD5, POLE1, RFC1 and RPA1; additionally PSF1 and PCNA1, which are downregulated. In HTT we found exclusively upregulated genes including ORC2, MCM3, MCM4, MCM5, MCM7, MCM10, TOPBP1 (MEI1), POLA3, POLA4, POLD1, POLD3, RFC2, RFC3, RFC4, RPA1, RPA2 and RPA3^25,51^. POLA2 lays down a short RNA/DNA primer in the lagging strand synthesis^25^, and is upregulated in both HPT and HTT datasets. Finally, the comparison of SeeTC.vs.HTT showed a shared upregulation of RPA2 with the HTT and a unique and strong downregulation for one RPA1 gene (Figure 6), both necessary to stabilize single stranded DNA. In summary, the HTT shows an upregulation of almost all the elements necessary for DNA replication, a characteristic behavior of cells going through endoreduplication (Figure 7).

**Figure 6.**
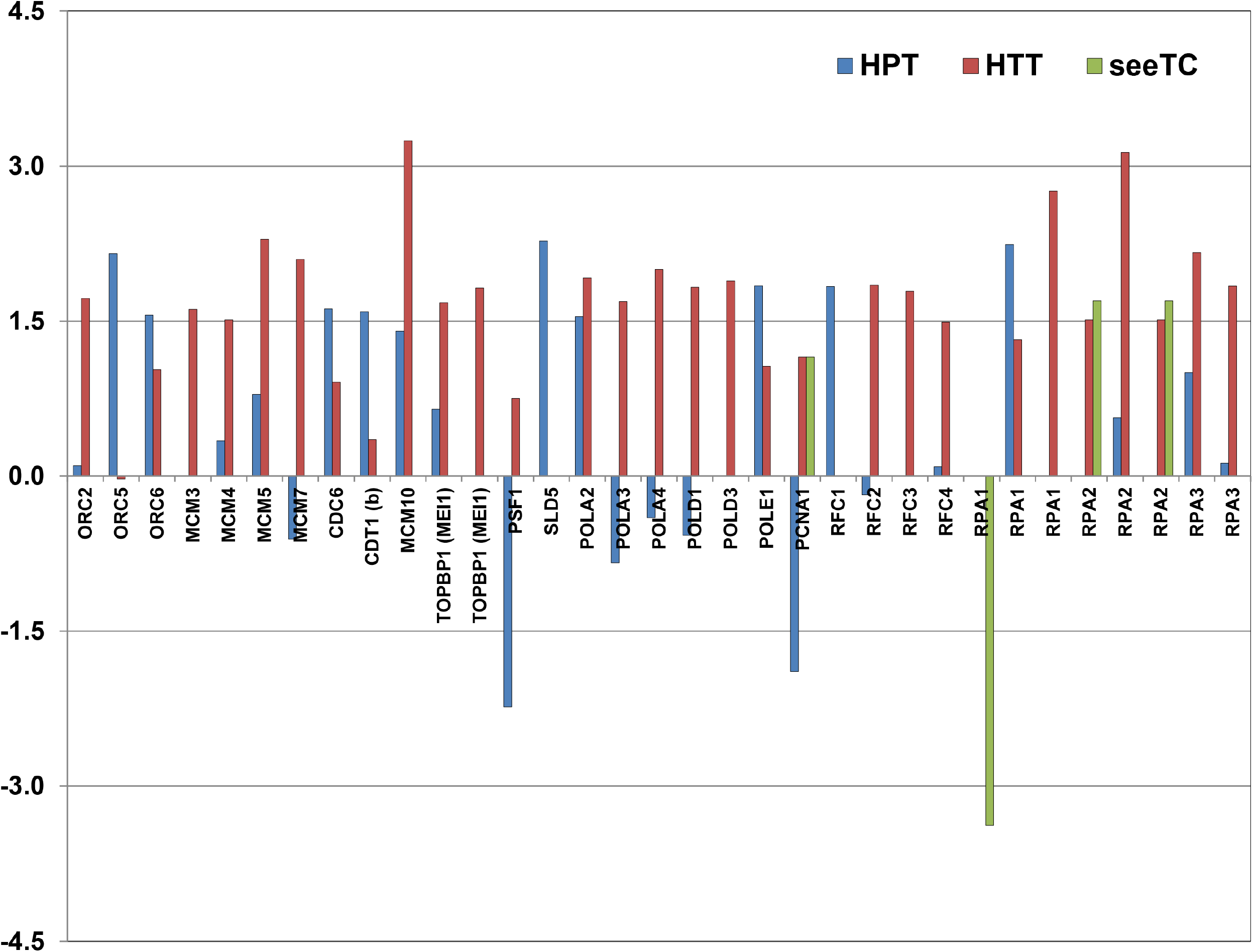
DE core DNA replication genes in response to *Ustilago maydis* infection in specific cell-types. Genes differentially regulated (FDR<5%) are shown. Y axis indicates Log2FC.

**Figure 7.**
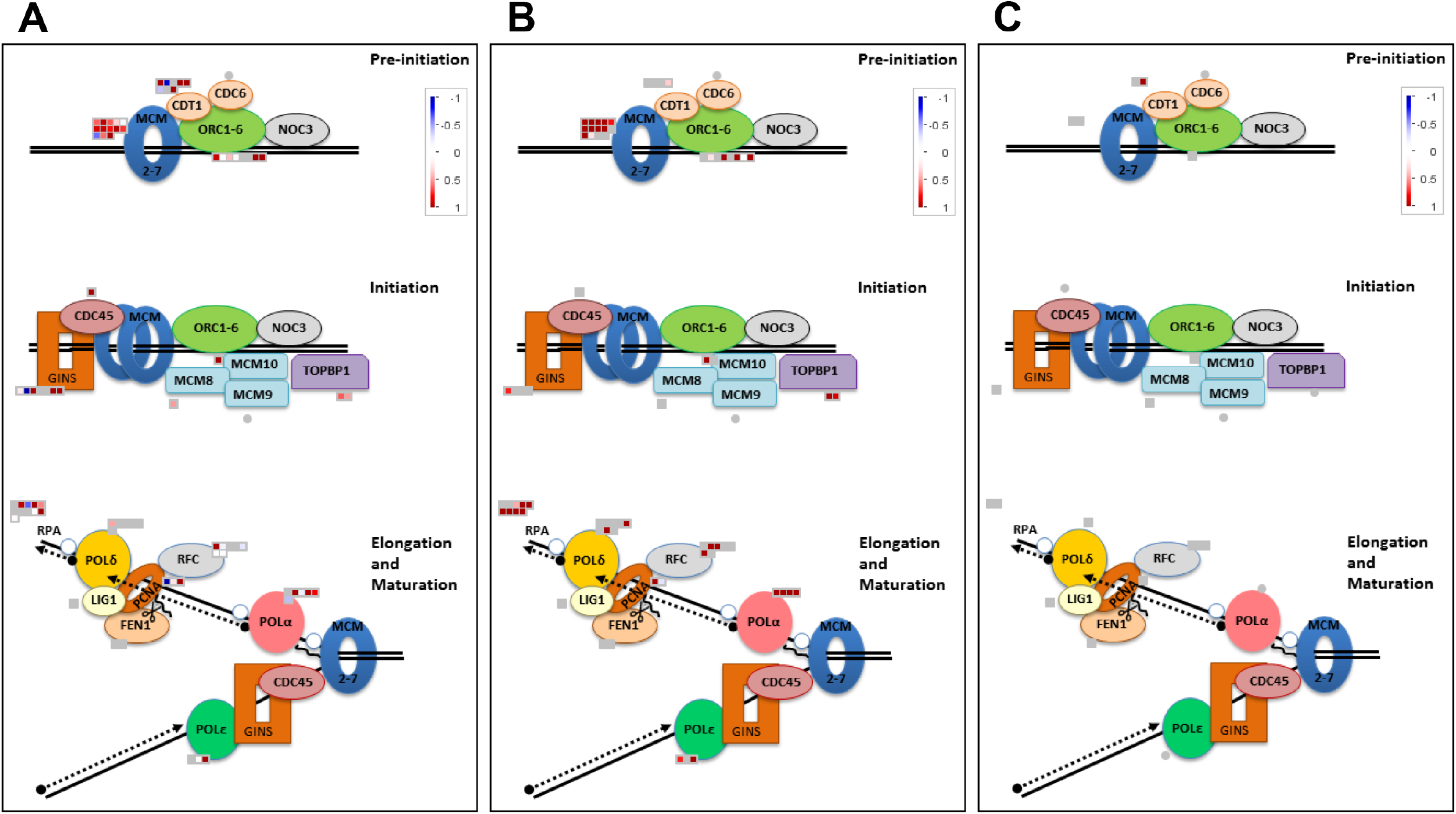
DE DNA replication machinery genes in response to *Ustilago maydis* infection in specific cell-types. **A** HPT. **B** HTT. **C** seeTC. Genes differentially regulated (FDR<5%) are shown. Upregulated transcripts are shown in red and downregulated transcripts are colored blue.

### The Skp1/Cullin1/F-box complex (SCF) and SGT1 interactors

The effector protein See1 is transferred from biotrophic *U. maydis* hyphae into the cytoplasm and, in particular to the nucleus of the host cell^8^. A yeast-2hybrid (Y2H) screen identified a maize homologue of SGT1 (suppressor of G2 allele of skp1) as its partner/target in maize^8^. SGT1 was originally identified as a cell cycle regulator necessary for the kinetochore formation in yeast^74^. It regulates the cell cycle together with Skp1 in two ways, by regulating Skp1 function in the Skp1/Cullin1/F-box complex (SCF), an ubiquitin ligase that controls the degradation of cell cycle regulators to allow G1-to-S transition, and by promoting the assembly of the centromere-binding complex that initiates kinetochore formation^74,75^.

Due to the important role of SCF in cell cycle regulation and the interaction of one of its subunits (SGT1) with See1, we decided to explore the expression of genes encoding for its components, additionally we included the 359 F-box genes reported by Jia et al., 2013^76^. F-box genes are crucial components of the SCF-ubiquitin ligases and confer substrate specificity, therefore, the higher the number of F-box proteins the more increases the number of potential SCF complexes. Our analysis showed DE genes encoding for SCF subunits in four out of five datasets (Figure 8). We observe upregulation of SGT1 in the HPT (Figure 8A). This observation might be relevant considering that See1 interacts with SGT1 and such interaction may have an impact on cell cycle as no hyperplasic cells are formed in maize leaves infected with SG200Δsee1^4^. In contrast, a general absence of SCF-complex activation is observed in SG200Δsee1 compared to SG200 infected mesophyll cells (Figure 8B and 8C).

**Figure 8.**
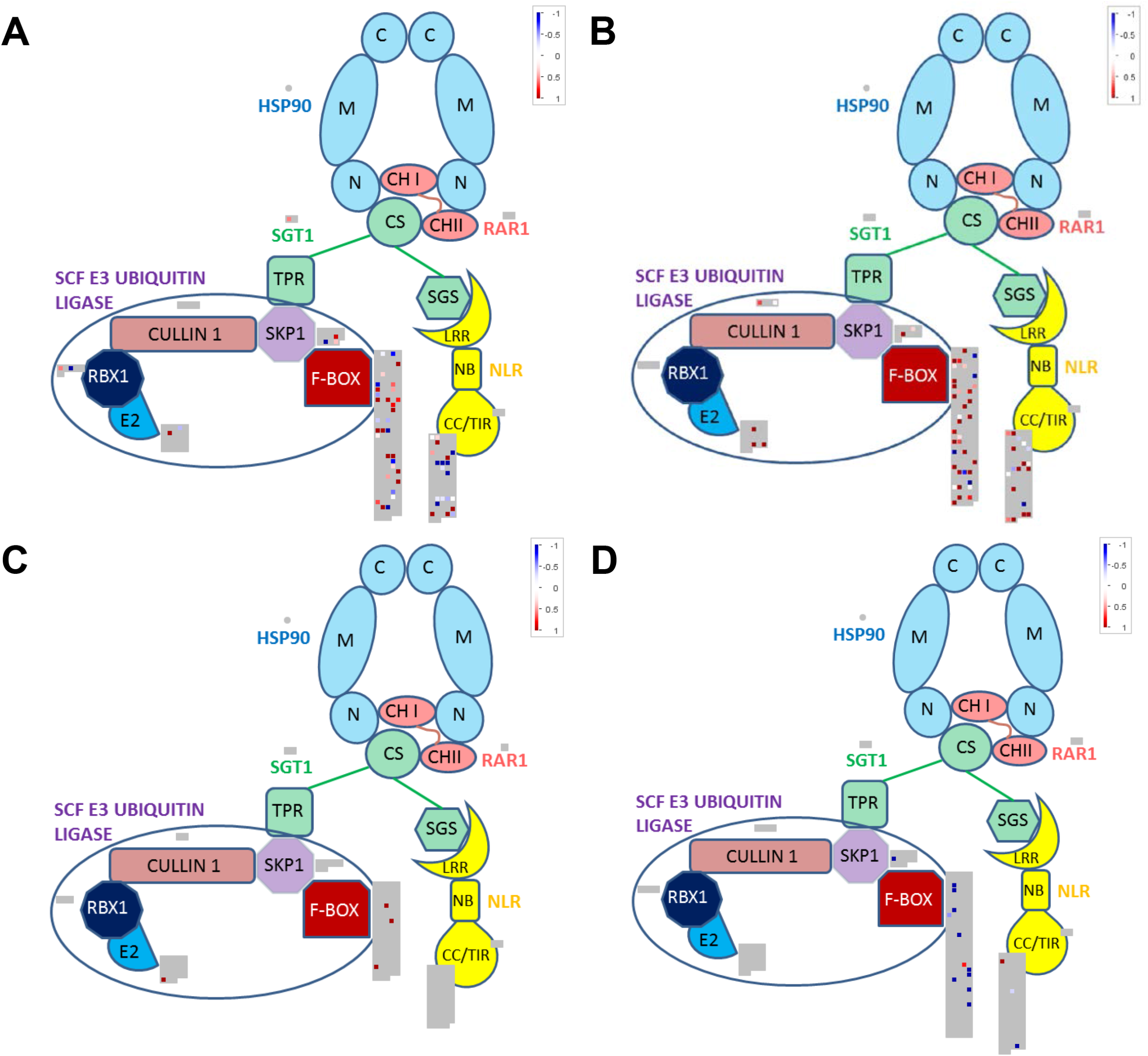
DE SCF subunit encoding genes in response to *Ustilago maydis* infection in specific cell-types. **A** HPT. **B** HTT. **C** seeTC. **D** SeeTC.vs.HTT. For the annotation of the genes encoding for the subunits HSP90, RAR1, SGT1, CULLIN1, RBX1, E2 and SKP1 the Mercator4 v1 program was used. F-box genes and NB-LRR-containing genes were based on the lists provided by Jia et al., 2013 and Song et al., 2015 with minimal modifications. Upregulated transcripts are shown in red and downregulated transcripts are colored blue.

8 F-box genes are upregulated in HPT and 11 F-box genes deregulated in the HTT dataset, from which two are strongly downregulated (Table III). In the seeTC dataset, 3 F-box genes were strongly upregulated (Table III). One DE F-box gene (ZmFBX154.1), upregulated in the HPT dataset, has been reported to respond to multiple stresses and may participate in the crosstalk between different signal transduction pathways^76^. The specific function of the majority of the F-box genes in plants remains unclear and only ZmFBX92, which is not DE in our datasets, has been functionally characterized^64,76^.

In summary, no strong expression changes in the SCF components were observed in the HPT and HTT datasets (Figure 8), but DE F-box genes were detected (Table III). Interestingly, only two F-box genes are both common and upregulated between the HPT and HTT datasets suggesting that the majority of the selective interactions of the SCF complex are specific for each tumor-type. As a consequence, the abundance of key regulatory proteins, among them proteins involved in the regulation of cell cycle, is likely specific for each tumor-type. We conclude that among the DE F-box genes strong candidates involved in the regulation of cell cycle can be found.

### The Small ubiquitin-like modifier (SUMO) and the SUMOylation machinery

SUMOylation is a post-translational modification that consists of the covalent attachment of a SUMO to a substrate protein^77^. SUMOylation regulates the activity of several proteins involved in critical cellular processes such as cell division and transcriptional regulation^77,78^. In yeast the SUMO-conjugating enzyme Ubc9 plays a role in the degradation of S and M-phase cyclins, and the ubiquitin-like specific protease ULP1, is essential for the G2 to M phase transition^79,80^. Furthermore, aberrant SUMOylation of key cell signaling proteins, including tumor suppressors and oncogenes result in deregulation of cell cycle and division, which ultimately leads to cancer^81^. In human cells, it was recently shown that SUMOylation of the APC4 subunit of the APC/C E3 ubiquitin-ligase is crucial for accurate progression of cells through mitosis; furthermore, SUMOylation increases APC/C ubiquitylation activity toward a subset of its targets^82,83^. In plants, SUMOylation has been implicated in several physiological responses and plays an important role to control cell cycle progression^84-87^. Particularly, the SUMO-E3-ligase AtMMS21 dissociates the E2Fa/DPa complex regulating in this way the G1/S cell cycle progression^88^.

Genes involved in the SUMO machinery are differentially expressed in both HPT and HTT (Supplementary Table 5). In the HPT dataset the only upregulated gene encodes for a SUMO conjugating enzyme subunit 1 (f) (ZmSCE1f, |log2FC| 2.59). In the HTT three SUMO machinery components are upregulated, a SUMO-variant (SUMO-V, |log2FC| 1.78), ZmSce1f (|log2FC| 1.76) and a SUMO ligase (SIZ1c, |log2FC| 2.63). From the three DE SUMO machinery members only ZmSce1f enzymatic function has been confirmed, while ZmSUMO-v and ZmSiz1c remain to be tested.

## Discussion

The full maize transcriptome analysis of SG200 infected mesophyll and bundle sheath cells has provided us a deeper view in the mechanisms evoked in the formation of maize leaf tumor. Expected responses, such as the alteration of genes involved in the regulation and performance of cell cycle, were differentially regulated in particular tumor cell types. Interestingly, some of the mechanisms observed differed between cell types and mostly reflected the cell behavior (i.e hyperplasic or hypertrophic). In comparison to the wealth of information and studies performed in mammals or yeast, plant cell cycle still requires a lot of study and homologues functionality validation. Such studies are complicated since in plants large families encode for cell cycle regulators^25,26^. It has been suggested that the evolution of larger families coding for CDKs and cyclins might help to provide a new layer of substrate recognition to coordinate the cell cycle with developmental cues^27^. More difficulties arise due to inconsistent nomenclatures, which difficult the comparisons and analysis^27^. A reductionist vision or description of the maize cell cycle would be simply not correct due to the lack of information/characterization of many genes. Most of the here reported genes still require a functional confirmation. However, this report gives some pointers to promising genes that could shed some light on cell cycles processes, i.e. endoreduplication. The identification of functional homologues that keep the network topology to control cell cycle will be crucial for the advance and understanding of this process in maize and other plants. Furthermore, previous comparative studies between plants and humans have identified putative cancer genes^89^. Such studies aim to identify conserved proliferation genes based on expression and transcriptional regulation in healthy tissues. Our study now provides data on cell cycle related genes in a tumorous tissue. Therefore we believe it provides promising candidates to understand tumorigenesis.

### Regulation of cell cycle and core-DNA replication machineries by *U. maydis* infection

In plants the analysis of cell cycle mutants has revealed that the loss of cell proliferation control is not sufficient to induce tumor development^24^. Furthermore, plants tolerate fluctuations in cell proliferation rates without this promoting tumor formation^24^.

Transcriptional activation of replication proteins (i.e. pre-replication complex) can induce endoreduplication^90^. Two *E2F* coding genes and one *DPA* gene are upregulated in HTT (Table II). The heterodimer E2F-DP promotes the expression of S-phase genes. Additionally, the majority of the components required for DNA replication are upregulated in the HTT dataset (Figures 5 and 6); this is in agreement with the hypertrophic phenotype observed^4^. Additionally, DPA expression levels have been also reported to correlate positively with final leaf size traits^64^. The RBR protein family is crucial and defined as a core cell cycle control by repressing G1/S phase cell cycle progression. RBR is known as a tumor suppressor and is inactivated in many human cancers^24^. Two *RBR* maize genes have been well characterized^66,67^, Zm*RBR1* has a canonical function as repressor of cell cycle progression while Zm*RBR3* promotes the expression of the E2F/DP targets, including the MCM family, required for the initiation of DNA synthesis^67^. Our analysis showed a strong upregulation of Zm*RBR4* and Zm*RBR3* in both tumor cell types but no alterations in Zm*RBR1/2* gene expressions (Table II and Figure 4). Zm*RBR4* has not yet been characterized but its strong expression in both tumor cell-types rather speaks for a positive role in cell cycle progression.

CKS2 is upregulated in the hypertrophic HTT cells. CKS2 is frequently overexpressed in human cancers and other malignancies and such overexpression overrides the intra– S-phase checkpoint that blocks DNA replication in response to replication stress ^91,92^, it is tempting to speculate that similar to human cancers, CKS2 upregulation allows DNA replication in despite the replication stress.

In eukaryotes there exists an overall similar topology controlling the entry into S-phase, while the control of mitosis through CDK phosphorylation-dephosphorylation cycles appears more diversified^27^. This reflects what we observe in our datasets where HTT “behaviour” fits with the predicted models while the hyperplasic cells or HPT, more dependent in rapid mitotic phases, is somehow more difficult to describe or predict based on the current observations, and therefore the pattern is more difficult to be described.

Endoreduplication can be achieved by elimination of mitosis promoting components in the presence of persistent DNA replication^90^. Several plant biotrophs induce localized host endoreduplication by activating common mechanisms that include the anaphase-promoting complex ad modulation of core cell cycle transcriptional machinery^93^.

Despite being a common mechanism in biotroph-plant interactions, little is known about the host proteins and mechanisms manipulated by the biotroph as well as the effectors involved^93^. The hypertrophic (HTT) cells present an upregulation of several D-type cyclins, E2F-DP, ZmRBR3/4 and the full pre-replication machinery, all necessary to support a persistent DNA replication. In this paper we shade some light on potential host protein candidates and the role of the *U. maydis* effector See1 in the stimulation of the endoreduplication process.

SGT1 is a protein that takes part in two important complexes, HSP90-RAR1-SGT1 and the SCF-E3 ubiquitin-ligase. HSP90-RAR1-SGT1 is essential in NLR-mediated immune responses and mostly localized in the cytoplasm^94^. On the other hand, SCF-E3 ubiquitin-ligase is crucial for the degradation of proteins involved in the regulation of cell cycle, and therefore mostly acting in the nucleus^75^. The upregulation of SGT1 in the HPT cells and the induction of hyperplasia in BS cells, a phenotype clearly absent in the maize leaves infected with SG200Δsee1, suggest that somehow the cell is reading out an absence of the SGT1 component, which could be due to “sequestration” via See1. It is tempting to speculate that See1 somehow fosters the localization of SGT1 into the nucleus, thus promoting the formation of the SCF-complex. Another possibility is that by occluding the phosphorylation site in SGT1^8^, See1 somehow fosters SGT1 interaction with SCF components instead of the HSP90-RAR1-SGT1 complex. This in addition would have on top the advantage of avoiding programmed cell death.

### SUMOylation machinery is induced in hyperplasic cells

In animal models, hyperplasia can result from the reactivation of pathways involved in embryonic development and suppression of terminal differentiation^95^. In humans, gathering evidence shows a close relationship between SUMOylation and cancer development, including progression and metastasis, with direct evidence that the deregulation of the SUMO-pathway affects the proper function of several oncogenes and tumor suppressor genes^96^. Furthermore, the SUMO machinery has been proposed as a cancer biomarker to determine malignant tissues and cancer progression^97^. Our transcriptome analysis suggests that, like in animals, the SUMO machinery members are specifically deregulated in oncogenic tissues.

ZmSCE1f is a representative isotype II of the SCE E2, this isotype is exclusively found in cereals^98^. Remarkably, all isotype II E2s were abundant in dividing tissues hinting a role during cell division^98^. We observe an upregulation of ZmSCE1f in both tumorous cell types, hypertrophic mesophyll cells (HTT) and hyperplasic bundle sheath cells (HPT); remarkably, such upregulation is stronger in the highly dividing hyperplasic cells, further supporting its role in cell division.

In maize, five SUMOs have been identified, three canonical SUMO genes including an identical duplication of SUMO1 (SUMO1a and SUMO1b) and SUMO2, an evolutionarily conserved SUMO variant (SUMO-v), and the cereal-specific DiSUMO-LIKE (DSUL)^98-100^. SUMO-v proteins are most closely related to the fungal/animal Rad60-Esc2-Nip45 (RENi) family, which is involved in DNA damage repair^101^. In maize, SUMO-v is expressed at moderate levels in all tissues, but little is known about its function^98^. Due to the conservation of interaction surfaces as the SUMO-Interacting Motif (SIM, which allows noncovalent interaction with SUMO) and β-grasp fold it has been suggested that SUMO-v may work as a recruiting partner or scaffold protein providing a surface for protein-protein interactions ^98^.

SIZ1c encodes for a SAP and MIZ/SP-RING type ligase and presents substantial sequence alterations affecting the PHD domain and C-terminal region, with minimal changes to the SAP and MIZ/SP-RING domains^98^. Since PHD domain is important for target recognition and ZmSIZ1c is highly expressed in the endosperm it is likely that such target substrates are endosperm specific^98^. ZmSiz1c is exclusively upregulated in HTT cells; whether similar target substrates normally expressed in the endosperm are awakening in the leaf tumor cells remain to be explored.

## Supporting information

Supplmentary Dataset 1. List of expressed maize genes

Supplementary Dataset 2. SEA comparison

Supplementary Dataset 3. SEA comparison uniques

## Additional Information

## Acknowledgments

This work was funded by the German Research Foundation (DFG; DO 1421/3-1) and the Cluster of excellence on Plant Science (CEPLAS).

## Author contributions

MV, AM and GD contributed conception and design of the study; MV, AH and CE performed the bioinformatics and statistical analysis; MV wrote the first draft; MV and GD wrote and edited the manuscript. All authors contributed to manuscript revision, read and approved the submitted version.

## Supporting Information

Additional Supporting Information may be found in the online version of this article.

**Figure S1** Chloroplast

**Figure S2** Starch – Sucrose

**Figure S3** Cell wall precursors

**Figure S4** Lignin synthesis

**Dataset S1.** List of expressed maize genes in all six datasets including estimated counts, fold-changes and adjusted p-values

**Dataset S2.** Singular Enrichment Analysis (SEA) and Parametric Analysis of Gene set Enrichment (PAGE)

**Dataset S3.** Singular Enrichment Analysis (SEA) and Parametric Analysis of Gene set Enrichment (PAGE) of unique genes

**Table S1.** GO enrichment terms with maximum percentage of genes shared between HPT and HTT datasets

**Table S2.** Unique GO enrichment terms in the HTT and seeTC datasets

**Table S3.** Core Cell Cycle Genes

**Table S2.** Core DNA Replication Machinery Genes

**Table S3.** SUMOylation components

