## Supplementary material for "Cell type specific transcriptional reprogramming of maize leaves during *Ustilago maydis* induced tumor formation": Supplentary Figures 1 - 4

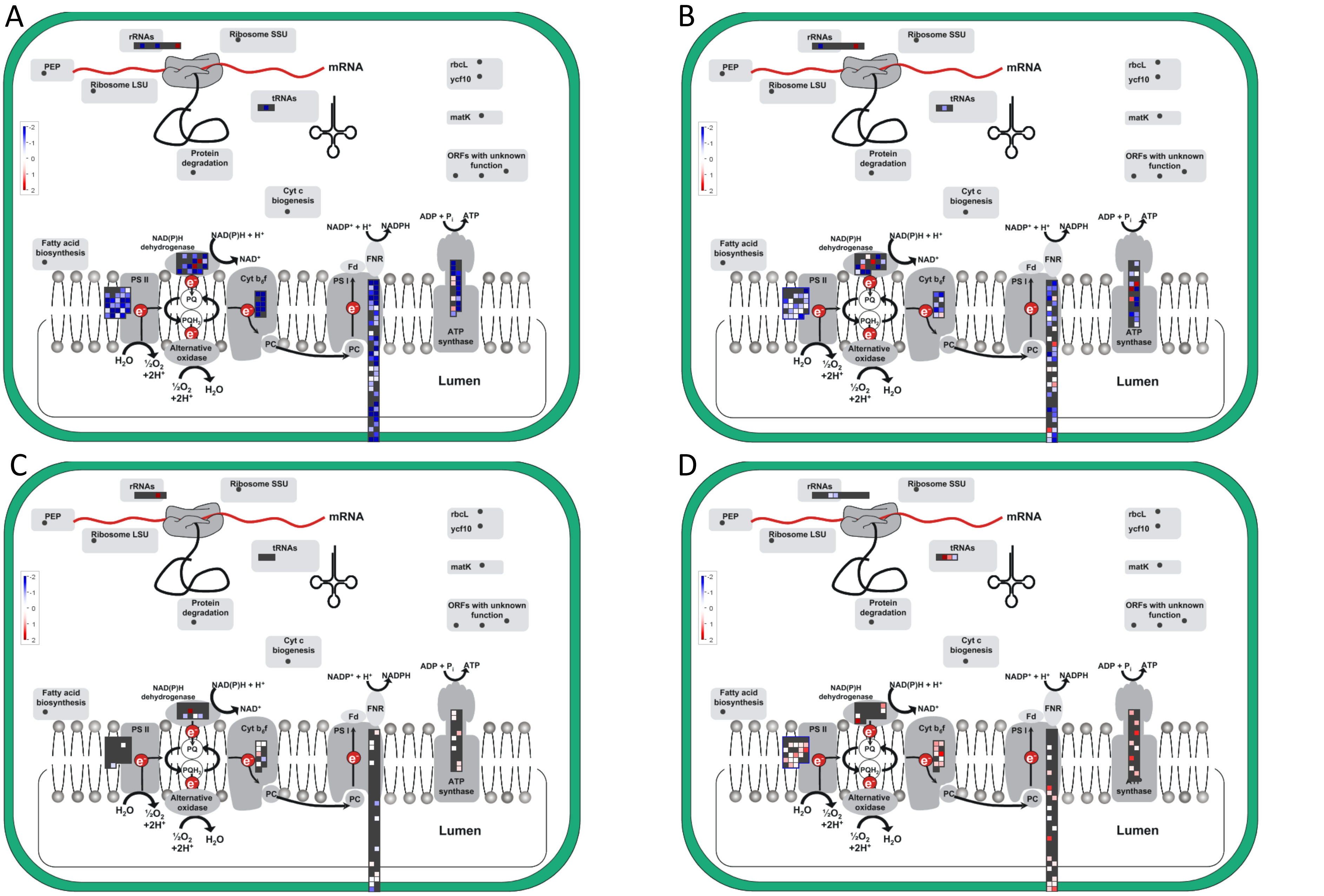


**Supplementary Figure 1.** Chloroplast responses to *Ustilago maydis* infection in specific cell-types. Genes differentially expressed (FDR ≤5%) are shown A, HPT. B HTT. C seeTC. D seeTC.vs.HTT: Upregulated transcripts are shown in red and downregulated transcripts are colored blue.

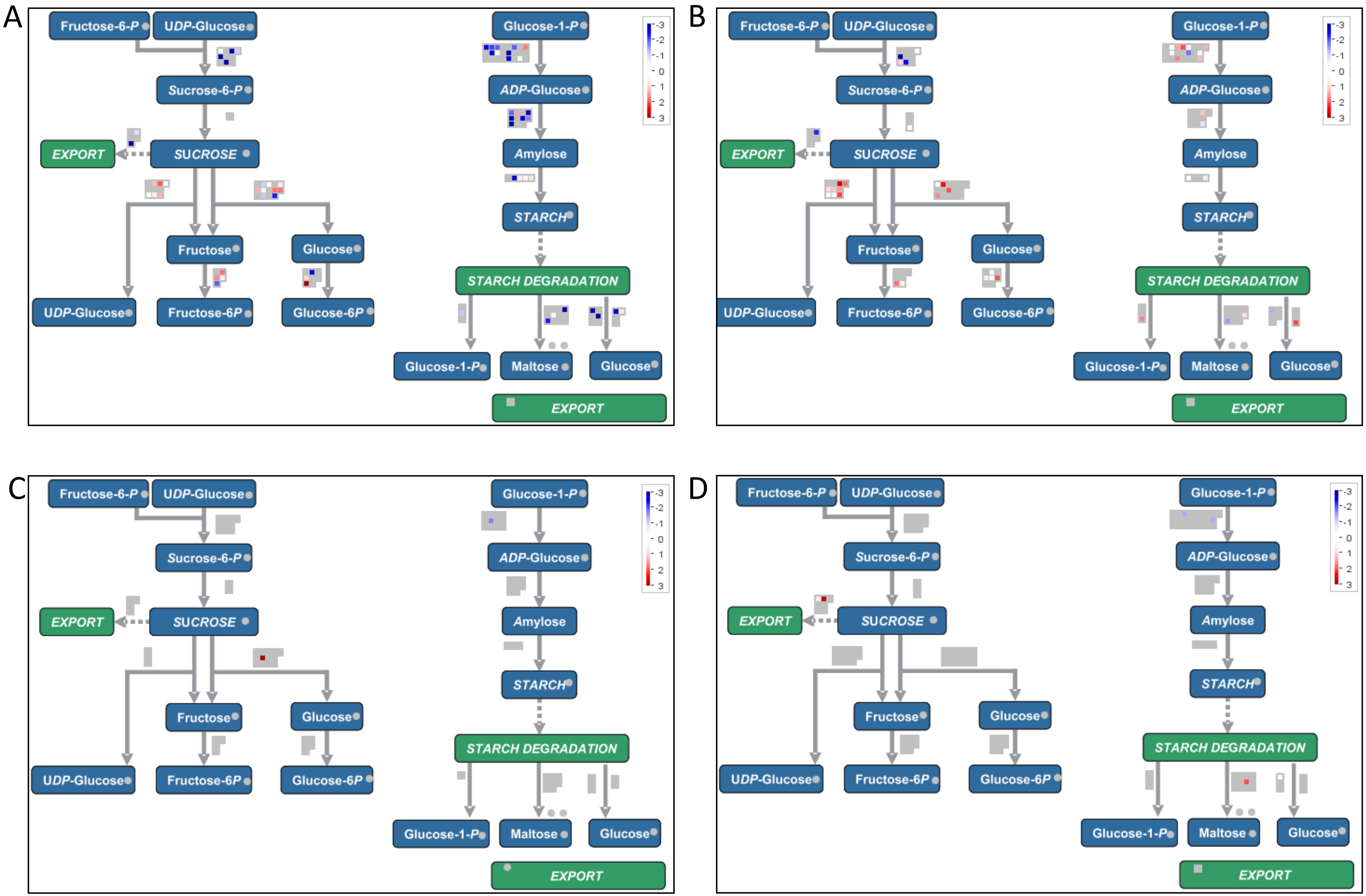


**Supplementary Figure 2.** Sucrose- and Starch- biosynthesis responses to *Ustilago maydis* infection in specific cell-types. Genes differentially expressed (FDR ≤5%) are shown A, HPT. B HTT. C seeTC. D seeTC.vs.HTT: Upregulated transcripts are shown in red and downregulated transcripts are colored blue.

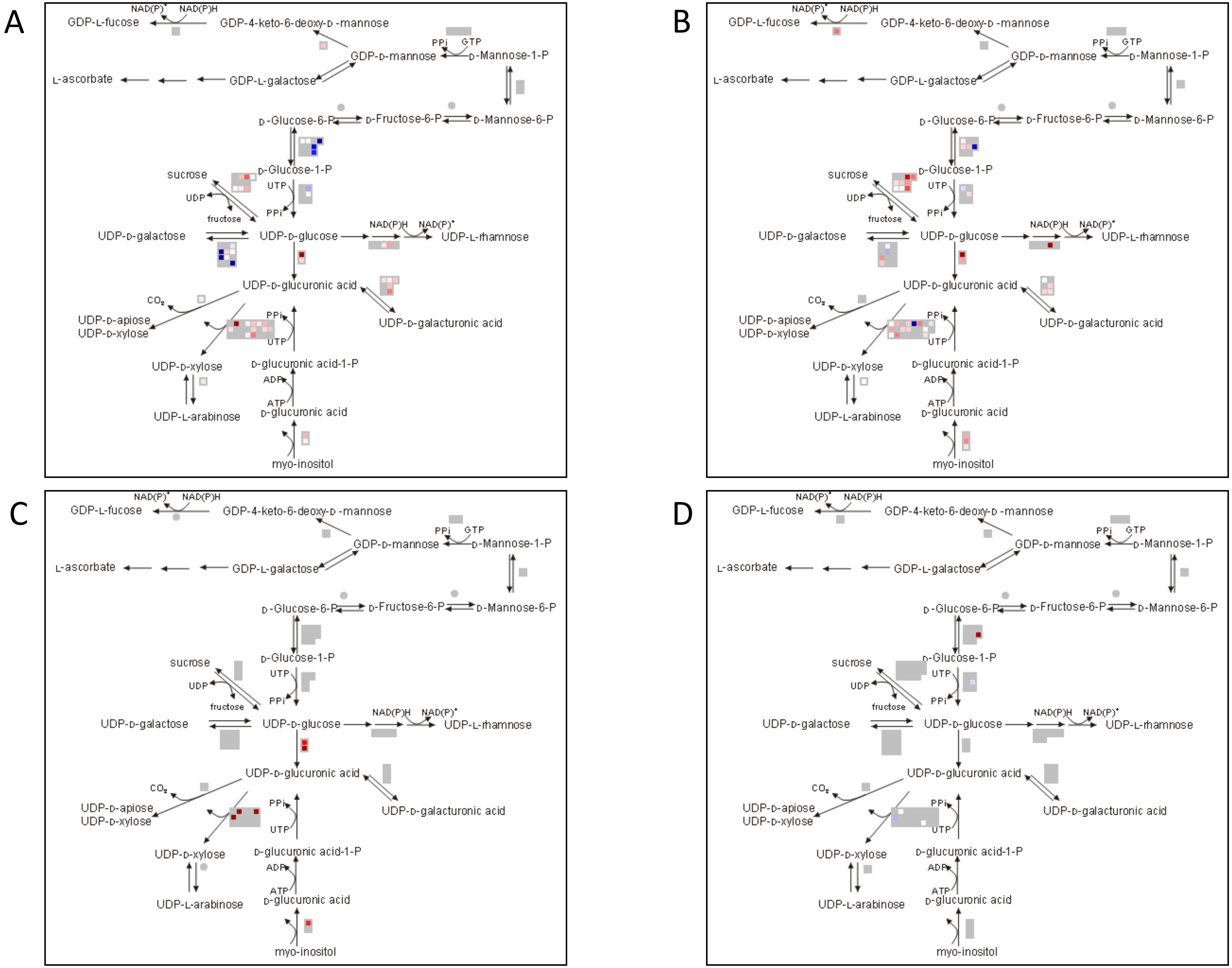


**Supplementary Figure 3.** Cell wall precursors biosynthesis responses to *Ustilago maydis* infection in specific cell-types. Genes differentially expressed (FDR ≤5%) are shown A, HPT. B HTT. C seeTC. D seeTC.vs.HTT: Upregulated transcripts are shown in red and downregulated transcripts are colored blue.

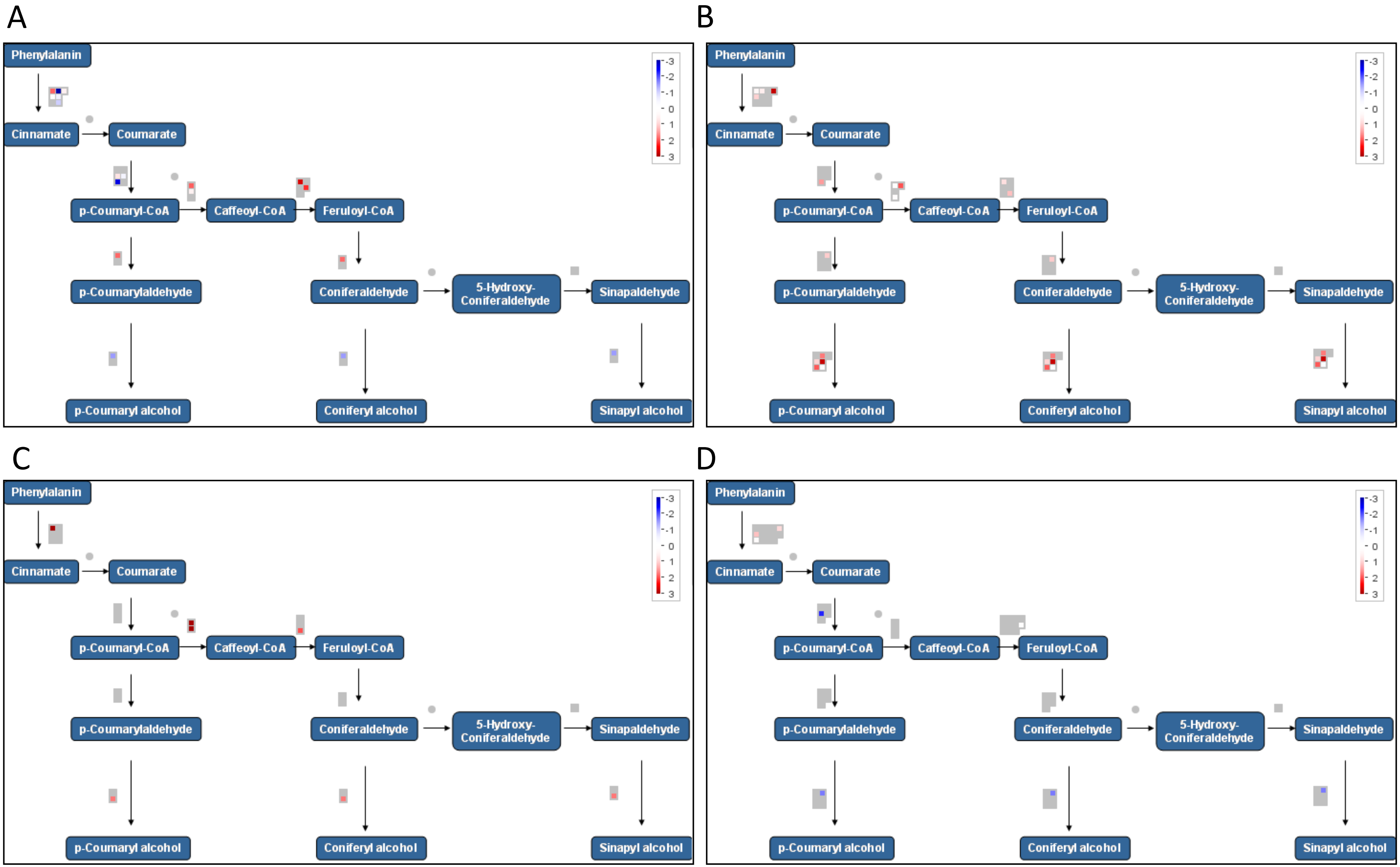


**Supplementary Figure 4.** Lignin biosynthesis responses to *Ustilago maydis* infection in specific cell-types. Genes differentially expressed (FDR ≤5%) are shown A, HPT. B HTT. C seeTC. D seeTC.vs.HTT: Upregulated transcripts are shown in red and downregulated transcripts are colored blue.
